# Invaders’ trophic position and their direct and indirect relationship influence on resident food webs

**DOI:** 10.1101/2025.01.23.634509

**Authors:** Ágnes Móréh, István Scheuring

## Abstract

Many ecosystems are undergoing simultaneous colonization and spread of multiple alien species. Invaders often negatively affect communities by reducing population sizes of resident species or even decreasing community diversity through extinctions. Direct and indirect interactions between them can amplify or mitigate their impacts on native communities. In this study, we compare the effects of two invaders on the resident model food webs under two scenarios: separate versus simultaneous invasion. We examined the resident food webs’ response from two perspectives: the number of extinct species as a measure of diversity loss, and the net change in total biomass of the food web. Using dynamic simulations based on the *Allometric Bioenergetic Model*, we tracked these changes *in silico* and compared the results of the two scenarios. We examined how the invaders’ trophic positions *relative to each other* and their direct/indirect interactions influence the additive or non-additive nature of the outcomes of their co-invasion. Our results have corroborated previous field and experimental observations, showing that when co-occurring invaders occupy different trophic levels, their combined effects may be dampened if one invader preys on the other. Further, we have shown that the probability of synergistic effect increases when invaders form a trophic cascade or share a common predator. However, we have also shown that all of these relationships are influenced by i) the metric used to track changes in the resident community, ii) the trophic level at which the invasion occurs, and iii) whether the invader has a predator (either resident or another invader).

**Highlights:**

- The impact of a joint invasion on the resident community depends on the used metric
- Additivity of the co-invasions’ impact is influenced by the invaders’ trophic position
- The effects of the invaders’ direct/indirect interactions depend on the trophic level
- Invaders in top position have stronger effects than those of intermediate position

## 1. INTRODUCTION

An increasing number of alien species are being documented globally, and numerous biological communities and ecosystems are facing, the challenges posed by multiple invasive species (Pyšek et al., 2020; Seebens et al., 2021). Resident communities are frequently subjected to simultaneous (or quasi simultaneous) invasion, and the interactions among the invaders can significantly influence community responses. These interactions often vary, making difficult to predict the overall effects on ecosystems (Crain et al., 2008; Darling and Côté, 2008). In our previous study (Móréh et al., 2024), we investigated how two invasive species influence each other’s success. Specifically, we examined how the positions of the invaders within the food web and the ecological interactions between them influence the success or failure of their joint invasion. However, non-native species not only affect each other, but more importantly from a conservation biological point of view, they also affect the invaded community (Guareschi et al., 2021; Preston et al., 2012). When an invasive species enters a food web and successfully establishes itself, it inevitably causes changes in the resident community, which can be tracked using various metrics (Móréh and Jordán, 2019; Pratt and Bowers, 1992). Naturally, to assess the magnitude of the impact, it is crucial to consider measurable attributes that allow for a comparison between pre- and post-invasion conditions, thus quantifying the effects of the invasion (Ricciardi et al., 2010).

The probability of whether an alien species will successfully establish a population in a novel environment depends on numerous factors, including environmental conditions (De Roy et al., 2013), resource availability (Kriksunov et al., 2011), compatibility with resident species, and the biological characteristics of the invader, such as its adaptability and ecological niche requirements (Vall-llosera et al., 2016). Additionally, this probability is often closely related to the number of individuals introduced (Cassey et al., 2018). Recent meta-analyses emphasized that the impact of invasive species on communities is closely tied to their abundance (Bradley et al., 2019) and trophic position (Thomsen et al., 2014).

When considering the impact of co-invasion on the resident community, it is inevitable to compare the combined impact of two or more non-native species with the sum of their impact when they invade individually. If the combined effect is equal to the sum of the individual effects, then invaders act additively. If the combined effect of the invasive species is smaller than the sum of individual effects, then co-invasion has an antagonistic effect, meaning it dampens the overall impact. If the combined effect of the invaders is larger than the sum of effects by separate invasions, then they interaction is considered as synergistic, meaning the effect on the resident community is amplified by co-invasion (Crain et al., 2008; Folt et al., 1999; Jackson, 2015; Preston et al., 2012). Numerous studies address two- or multi-species invasions, incorporating both field and mesocosm experiments (Braga et al., 2020; Figueroa-Muñoz et al., 2022; Gao et al., 2021; Johnson et al., 2009) alongside meta-analyses that synthesize empirical observations (Cameron et al., 2016; Jackson, 2015). However, the effects of invasive species on native communities arises from complex ecological interactions that make it challenging to quantify and predict their impact through empirical studies. Consequently, theoretical models can play a special role in this field, helping to explain the field and experimental results, and providing insights for new research directions. Surprisingly, such models are relatively scarce. Hewitt and Huxel (2002) explored colonization success using the standard Lotka-Volterra assembly model. Their findings revealed that inoculation density had a more significant impact on invasion success than the number of simultaneous invaders, as evidenced by the increased rate of successful invasions. Rauschert and Shea (2017) applied a spatially explicit competition model to examine invasional interference between two invaders. Their study focused on how the strength of inter- and intraspecific interactions influence the coexistence of the invaders. In natural food webs, co-occurring invaders often occupy different trophic positions, and their complex interactions play a crucial role in determining overall invasion outcomes (Preston et al., 2012). Other previous studies demonstrated that factors reflecting the interaction between the invader and the resident community, such as trophic position or trophic generality, are key predictors of invasion success (Jones and Gomulkiewicz, 2012; Romanuk et al., 2009). Another crucial aspect of understanding the impacts of alien species is exploring both their direct and indirect interactions, such as predation (Penk et al., 2017), trophic cascade (Liversage et al., 2021) or competition (Preisser and Elkinton, 2008; Ross et al., 2004). Vagnon et al. (2022) applied an allometric niche model to infer the trophic characteristics of two invaders in a French lake. Their study highlights the potential impacts of this multi-species invasion on this ecosystem, particularly through alterations of trophic links within the invaded food web. Here, we use an allometric model to study food webs, focusing on whether differences or similarities in trophic position, trophic level, and direct/indirect relationships between two invaders in simultaneous invasions influence the additivity of their impacts. We aim to address the following questions:

- How does the trophic position of two invasive species affect the impact of their joint invasion on the resident food web? Specifically, how does the trophic level of the invaders, either individually or in combination, influence the extent and nature of changes in species diversity and total biomass of the food web?
- Does the topological distance between two invaders within the food web, and their ecological relationship (e.g., predator-prey, exploitative competition, or trophic cascade) affect the frequency of additive/non-additive outcomes? Furthermore, how do these effects differ across various trophic levels?

## 2. METHODS

In this study, we examine the magnitude of the combined effect when two invasive species appear in the resident food web simultaneously and compare this to the scenario where they invade separately. Our study focuses on artificially generated communities, which simplify their handling and comparison by allowing many network properties to be fixed. This approach limits the scope of measurable traits, as we only record a few, more generalized characteristics of the resident networks during simulations:

i. *Change in species diversity of resident food web* (Fig. 1), where we compare the number of species extinctions caused by each invader individually to the total number of extinctions resulting from their joint invasion (hereafter refers to as EXT index). The effects of the two invaders can overlap (Jackson et al., 2016; Tekiela and Barney, 2017), meaning that both invaders may cause the extinction of some or all the same species, so we define additive, antagonistic and synergistic effects by considering this fact. The outcome is additive if the union of the extinctions caused by the two invaders acting separately is identical to the extinctions caused by the joint invasion. Joint invasion is antagonistic (synergistic) if the union of the number of extinct species is smaller (bigger) than the union of number of extinct species as a result of separate invasions (Fig. 1). For example, if the first invading species (I1) causes the extinction of 2 species and the second invading species (I2) also causes the extinction of 2 species, but 1 extinct species is common to both, the total number of unique extinctions across the two invaders would be 3, so the invaders are in antagonistic interactions.
ii. *Change in total biomass* of the resident food web (Fig. 2): we compare the total biomass of the post-invasion community in case of individual and joint invasions too, (ignoring the invaders’ biomass) to the baseline biomass of the native food web prior to invasion (hereafter referred to as BM index). To improve comparability of results, we measure relative changes. In this case, beyond the outcomes defined above (additive, antagonistic or synergistic), the biomass change observed during joint invasion can sometimes occur in the opposite direction compared to the changes caused by each invader separately (Jackson et al., 2016; Piggott et al., 2015). This “reversed” effect represents a distinct subset of antagonistic outcomes (Fig. 2). If the two separate invasions cause changes in opposite directions in the resident food web (e.g., one increases and the other decreases the biomass), the joint effect is categorized as reversed if it surpasses, in magnitude, the larger of the two separate effects while acting in the opposite direction (see Fig. 2b). The antagonistic, additive and synergistic categories just defined will now be referred as additivity categories from now for the EXT and BM indices.

**Figure 1.**
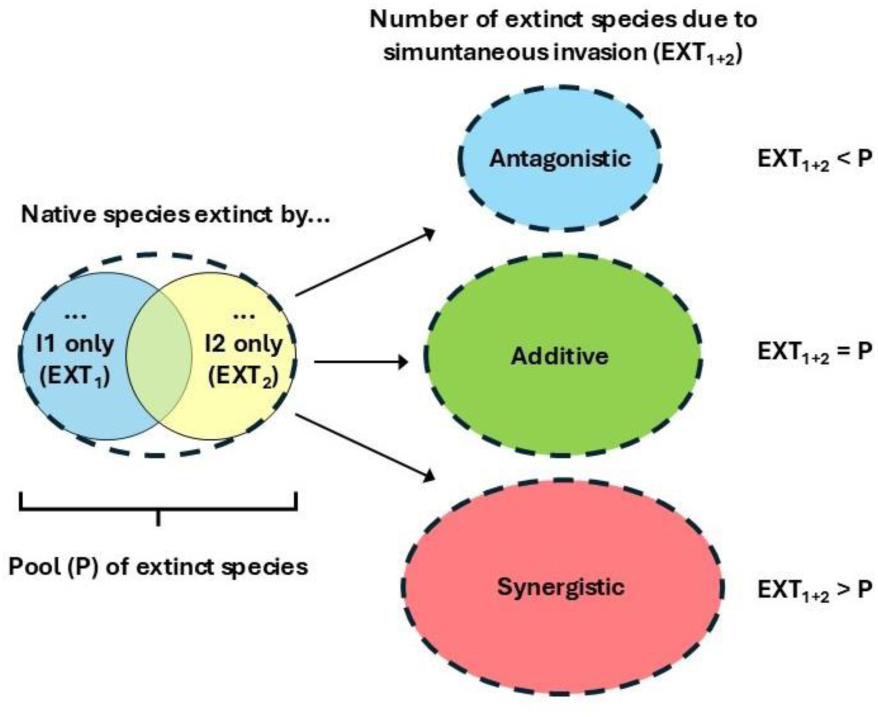
Schematic representation of the joint effect of invaders when considering the decline of biodiversity (EXT index): the combined pool (P) of species extinct by each of the two invaders separately (I_1_ and I_2_) serves as the baseline to assess the cumulative impact of the invaders (EXT_1+2_) when they invade together.

**Figure 2.**
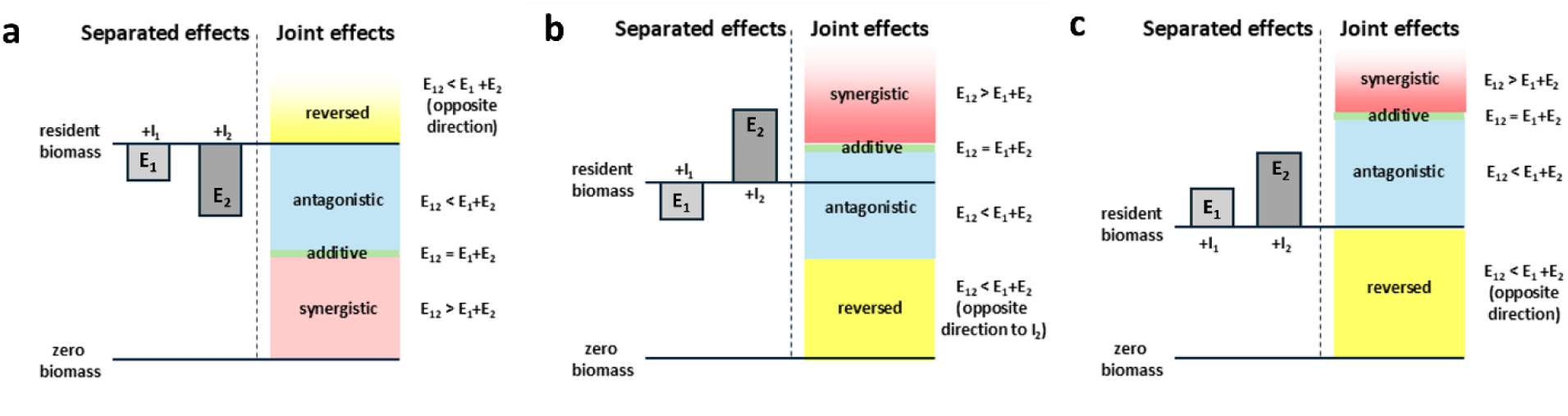
Schematic representation of the additive and non-additive effects when tracking the biomass change of the resident food webs caused by the invaders’ separate (E_1_ and E_2_) and joint invasion (E_12_).

### Construction of resident food webs and simulation of invasion

The method and steps for both the generation of resident food webs and the simulation of paired and separate invasions into them are the same as described in our previous work (Móréh et al., 2024), so we only give a brief overview here.

We employed the *Allometric Bioenergetic Model (ABM)*, an allometric coefficient-updated consumer-resource model, for both resident network generation and invasion simulation (Brown et al., 2004; Yodzis and Innes, 1992). This framework is particularly effective for modeling various types of food webs, especially aquatic communities. Changes in species biomass density (*B_i_*) are described by a system of differential equations:

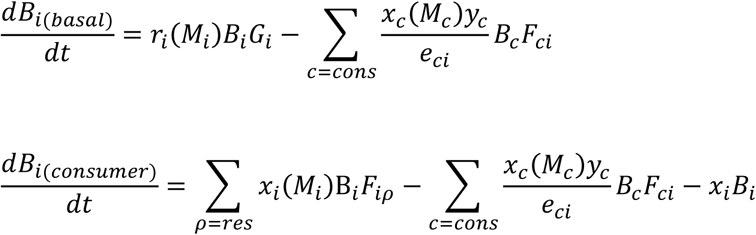

The first equation applies to the basal species, where the first term on the right-hand side represents the growth of the basal species. The mass-specific maximum growth rate of species *i* is represented by *r_i_(M_i_)*, and *G_i_* is the term for the logistic net growth rate. The second term expresses the biomass-decrease of the basal species due to consumption: *x_c_* represents the consumers’ mass-specific metabolic rate, and *y_c_* denotes the maximum consumption rate relative to its metabolic rate. *F_ci_* is the nonlinear functional response term that represents the actual consumption rate when species *c* consumes species *i*, and *e_ci_* represents species *c* assimilation efficiency when consuming species *i*. A detailed explanation of the terms *G_i_* and *F_ij_*, as well as the mass dependence of each parameter and the allometric constants used, can be found in section S1. of Supplementary Material 1 (SM1).

Resident communities of different sizes (10-20 nodes) are generated by considering several topological rules (Dunne et al., 2002a; Hall and Raffaelli, 1991; Williams and Martinez, 2004). The number of basal and top species can vary from 1 to 4, with a maximum of five trophic levels. Omnivory is not excluded, but loops are, i.e. each species can only feed on a trophic level lower than its own. Since network connectance affects invasion success (Baiser et al., 2010; Romanuk et al., 2009), its value is fixed (*C*=0.2). The connectivity value and the number of nodes (*S*) determine the number of links (*C*=*L^2^S*). The maximum number of links per node was determined following Dunne et al. (2002b) (2*L*/*S*). A food web was considered suitable for invasion analysis if i) it had a stable fixed point in the steady state, and ii) all the resident species were co-occurring.

To simulate invasion, we extended the resident food webs by adding the invading species, depending on the specific scenario studied (separate or joint invasion). The invader pairs were systematically distributed across the consumer spectrum of the network (for technical details, see section S2. in SM1). If a drawn combination reached a stable state by the end of the simulation, it was kept for further analysis. This process resulted in a total of 121.000 invasion scenarios with 110 resident food webs (10 of each size) available for testing. In this study, we focused on comparing the combined effect of invaders on the resident food web to their individual effects. To ensure consistency, we only analyzed invasion scenarios where both invaders successfully established stable populations in both separate and joint invasion cases. This approach narrowed our analysis to the impact of 66,040 successful invasion events (see Móréh et al., 2024), excluding cases where at least one invader failed to establish in either scenario.

### Influencing factors

To determine the invades’ trophic position, we categorize the invaders first based on their feeding type (Preston et al., 2012) into the following groups:

- Primary consumer (*I*) - feeds exclusively on basal species.
- Omnivore (*O*) - feeds on both basal species and primary consumers.
- Secondary consumer (*II*) - feeds exclusively on primary consumers.

Within each of these categories, we also examine whether the species occupies a *top* or *intermediate* position within the food web—that is, whether it has a predator or acts as an apex predator. However, it is important to determine whether a predator-prey relationship exists between the two invaders. While each invader might independently occupy a top position, their joint appearance could result in one becoming a predator and the other its prey, thereby shifting one invader to an intermediate position.

To quantify the positions of invaders within the resident food web, several network indices can be calculated to reflect their structural and positional significance (Jordán et al., 2007; Wasserman, S. and Faust, K., 1994). Among these indices, the *trophic level* (TL) index is most directly related to the dietary levels described above. In this study, we used the prey-averaged trophic level metric, which is calculated as 1 plus the average trophic level of all the prey species consumed by a given consumer (Levine, 1980; Williams and Martinez, 2004). As in our previous study, we examined whether the *difference* (i.e., the extent of variation between values) and the *average* (i.e., how high or low the values are on average) of the invaders’ TL-indices influence the outcome of a joint invasion—in this case, the level of additivity. These two approaches are complementary: a small difference can occur with either low or high values, while a medium average can correspond to either high or low differences. To ensure comparability of the results across different food webs, the TL indices of the invaders were normalized to a scale between 0 and 1, and the results are reported using these transformed values.

Another important factor is the type of the ecological relationship that develops between the two invaders during the joint invasion (White et al., 2006). Do they have a predator-prey relationship, share a prey or predator, or participate in a trophic cascade? These interactions can significantly influence the dynamics and outcomes of their co-invasion (Preston et al., 2012). In this study, we examined the same categories as in our previous work (Móréh et al., 2024), their schematic representation and frequency of occurrence are shown in Fig 3. The three main categories of invader relationships are:

1. *1 step (close) connection:* direct predator-prey relationship exists between the invaders.
2. *2 steps apart:* trophic cascade or competitive relationship occurs between the invaders.
3. *Distant connection (>2 steps apart):* the invaders are separated by a greater topological distance in the food web.

**Figure 3:**
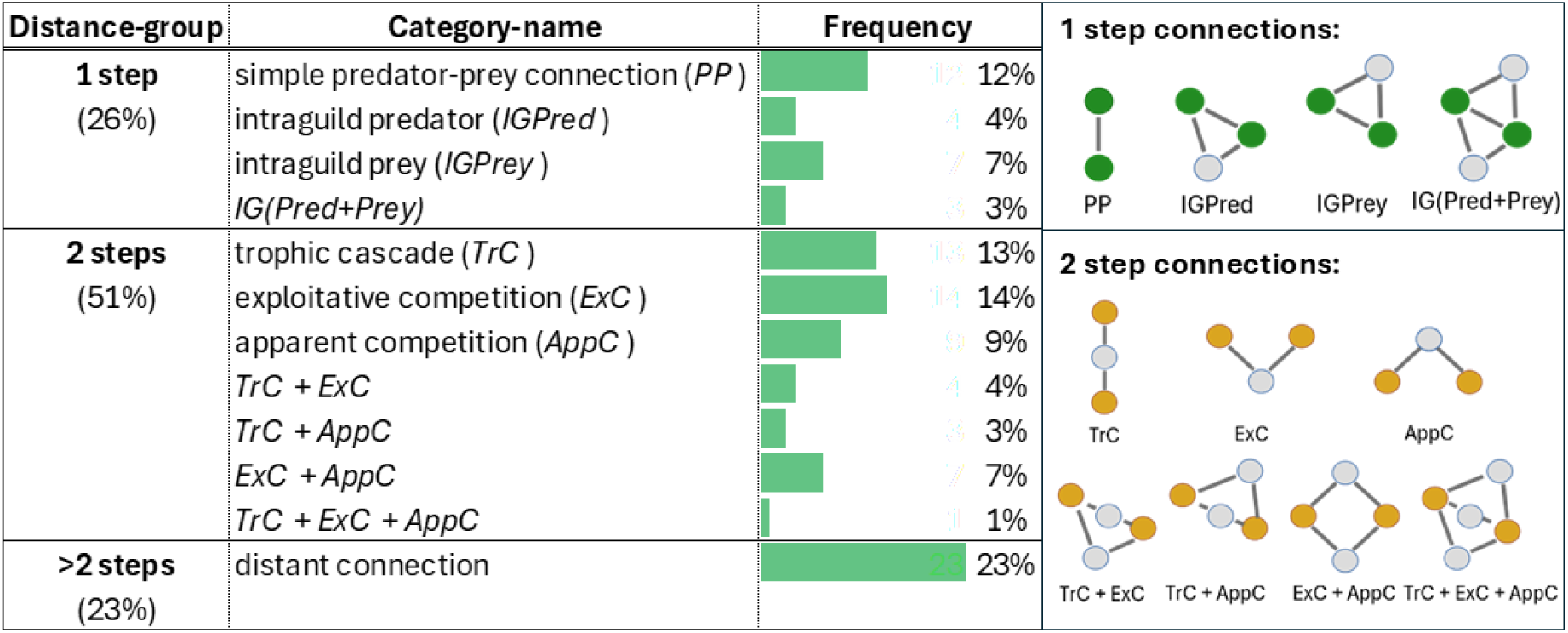
Frequencies and schematic representations of the investigated categories of the invaders’ ecological relationships of the successful invasion events. Invaders are represented with green (1 step connections) and orange (2 step connections) circles.

Naturally, these relationships are influenced by the trophic positions of the invaders—species at very different trophic levels are more likely (but not necessarily) to have a greater topological distance between them.

The generation of the resident networks and the simulation of invasions were carried out using a custom C code with the CVODE differential equation solver (Hindmarsh et al., 2005). The evaluation of the results, statistical analyses, and visualizations were performed in R (R Core Team, 2022). A summary of the statistical analyses used is provided in section S3. in SM1., and detailed results of the tests can be found in the tables of Supplementary Material 2.

## 3. RESULTS

### 3.1 Frequencies of the different additivity categories

The distribution of additivity categories differed notably between the two tracked indices. For the EXT index (Fig. 4a), roughly one-third of the cases (35%) showed no extinctions due to invasion, meaning that the resident food web remained intact in both the separate and simultaneous invasion scenarios, even though the invaders successfully integrated into the food web. Additive outcomes were observed in 28% of cases, while antagonistic and synergistic outcomes occurred in 21% and 16% of cases, respectively. For the BM index (Fig. 4b), successful invasions always led to changes in the overall biomass, even if they did not result in the extinction of any species. In most cases (60%), the combined effect of the two invaders was antagonistic, while 26% resulted in synergistic outcomes. Additive effects were relatively rare, occurring in only 10% of cases, and reversed effects were observed in just 4%.

**Figure 4.**
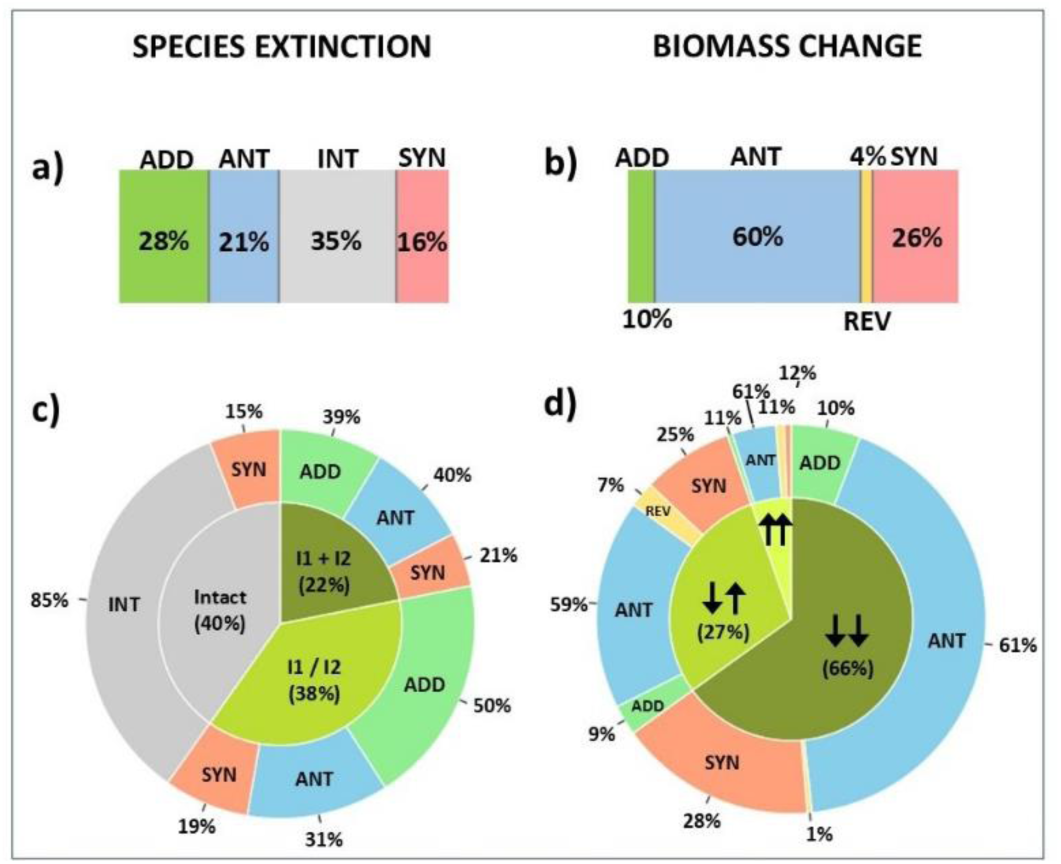
Frequencies of the different additivity categories a-b) Overall frequencies of the different additivity-categories in case of the two investigated indices (ADD: additive, ANT: antagonistic, SYN: synergistic, INT: intact, REV: reversed); c-d) Inner pie chart: the frequency of subgroups, categorized by the presence (both invaders (I1+I2), only one invader (I1/I2) or neither invader (Intact) cause extinction) and direction of the effects of the invaders (arrows) on the resident network; Outer donut chart: the distribution of additivity categories within these subgroups, provide insights into how different invasion scenarios influence the resident network.

The frequency of outcomes is influenced by whether or not each invader causes extinctions and by the direction of their effect on the resident food web biomass, either reducing or increasing it. For the EXT index (Fig. 4c), in 22% of the cases, both invaders caused extinction (I1+I2, inner chart), where the frequency of additive and antagonistic cases is almost equal (39% and 40%). If only one of them (I1/I2, inner chart) causes extinction separately (38%), then the probability of joint invasion causes additive outcome not surprisingly is significantly higher (50%). If the invaders have no effect separately on the species composition of the web, they are highly likely to have no effect together too (85%), but they may still cause extinction together, in which case their effect is naturally synergistic (15%).

For the BM index (Fig. 4d), both invaders predominantly reduce the biomass of the resident food web (66%), while they have opposite effect separately in 27%, but in some cases, and they even can increase the total biomass. Antagonistic outcomes predominate in all subcases (61%, 59%, 61%, outer chart in Fig. 4d). Similarly, the proportion of additive cases does not vary significantly but covers a much less fraction from the cases (10%, 9%, 11%, outer chart in Fig. 4d). The probability of synergistic outcomes is highest (28%) when both invaders reduce biomass, while the probability of reversed cases increases when at least one of them has a biomass-increasing effect (7% and 11%, outer chart In Fig. 4d).

### 3.2 Impact of the invaders’ trophic position

The lower the trophic position at which an invader enters the food web, the more likely it is to negatively impact the community by reducing diversity and biomass (section S4. of SM1.). Additionally, whether the invader has a predator (either native or another invasive species) or occupies a top position is crucial. In the latter case, the invader tends to exert a greater impact on the network (section S4. of SM1). These factors also influence the probability of additive or non-additive outcomes.

We analyzed how the frequencies of each additivity category changed with the trophic positions of the two invaders expressed by their feeding type (primary/omnivore/secondary consumer), comparing these values to the overall sample frequencies (Fig. 4a and 4b). Detailed results of the χ^2^ tests and the post hoc z tests can be found in Tables S1. For the EXT index, there is a strong significant relationship between the feeding type of the invaders (primary/omnivore/secondary consumer) and the additivity of their joint impact on the community (χ^2^(15, *N*=66040) = 7353, *p* < 0.001, Cramer’s V = 0.19, Fig. 5a). Note, that due to the high degrees of freedom, even a lower Cramer’s V value can indicate a relatively strong relationship (Cohen, 1988). The frequency of additive outcomes varies less with the dietary types (Fig. 5a,b). When both invaders are primary consumers (I+I), the frequency of synergistic outcome is highly increased (48.8%) comparing to the overall probability (17.5%), while it is reduced at the other trophic positions in the same comparison (omnivore or secondary predator) (Fig. 5a). However, the probability that the resident network food web remains intact (i.e., no extinctions occur) steadily increases as the invaders shift from being primary consumers to secondary consumers. In parallel, the probability of a synergistic outcome steadily decreases (Fig. 5a).

**Figure 5.**
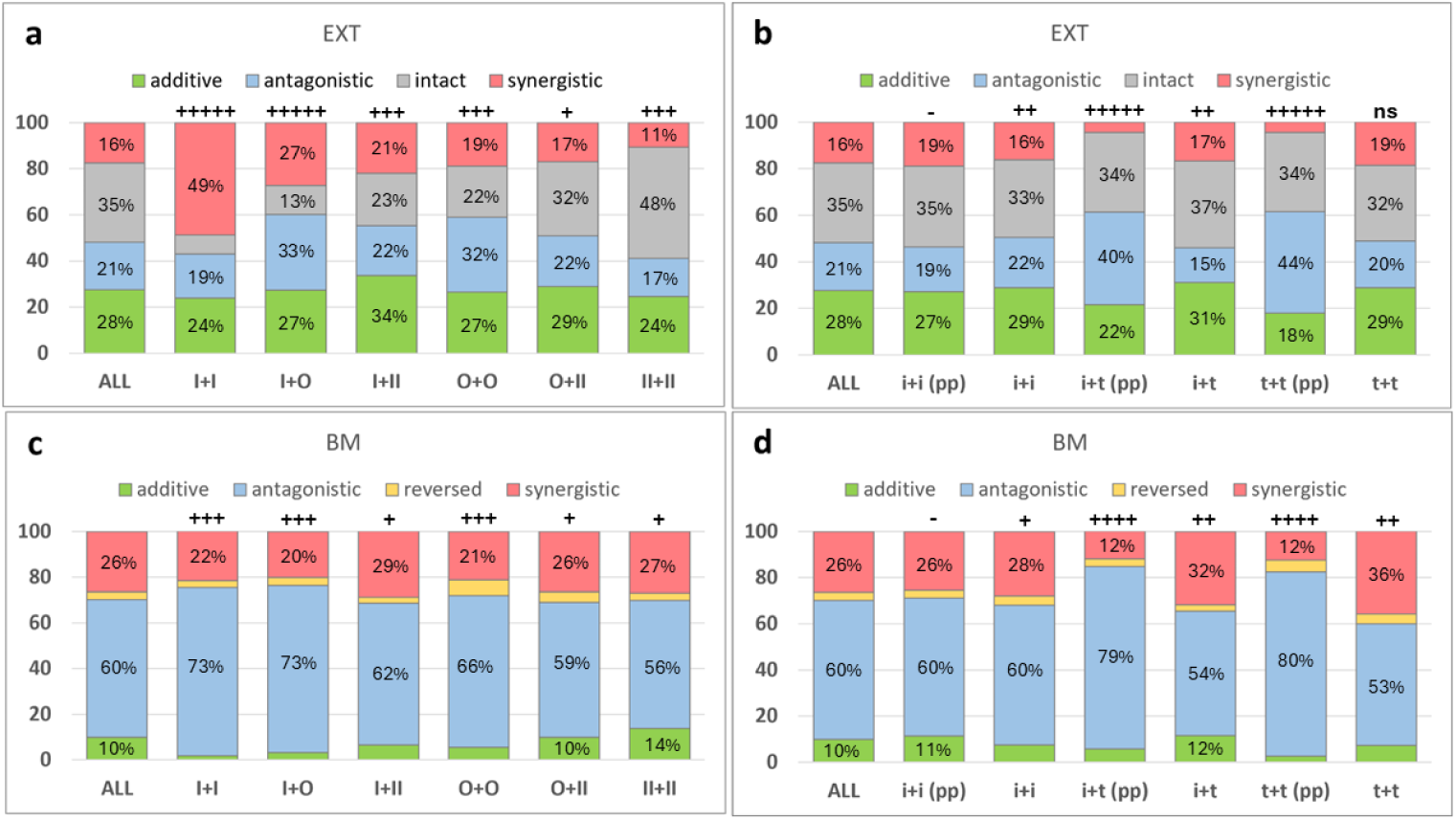
Frequency distributions of the different additivity categories are shown in function whether the invaders are primary consumers (I), omnivores (O) or secondary consumers (II) (a,c), and, in function of their intermediate (i) or top (t) positions (b, d). pp refers to the situation when the invaders form a predator-prey connection when invading simultaneously. The first columns on the plots show the frequency distribution in the whole dataset (ALL), while the symbols above the rest of the columns indicate whether the effect size (Cohen’s ω) is negligible (-), small (+), moderate (++) or medium (+++) or large (++++), indicating the magnitude of the deviation from the full sample. ns means that the difference is not statistically significant. Frequency-values below 10 % are not displayed on the bars.

Whether the invaders act as an intermediate (*i*) or top (*t*) consumer is only weakly related to the type of joint effect overall (χ^2^(15, *N* = 66040) = 919, *p* < 0.001, Cramer’s V = 0.07). However, the paired proportions test clearly shows that while the intermediate/top position in itself has small influence on the frequency of the different additivity categories, if at least one of them is a top predator (i+t and t+t), the possible predator-prey relationship between them significantly increases the likelihood of antagonistic effects (see Fig. 5b). Interestingly, this increase does not occur when both invaders are in intermediate positions (i+i).

For the BM index, although significant, we see only a weak relationship between the frequency of each category and the type of invader diet (χ^2^(15, *N*=66040) = 1653.1, *p* < 0.001, Cramer’s V = 0.09), with a weak trend: the higher the trophic status of the two invaders, the higher is the likelihood of an additive effect, while the lower positions increase the probability of more antagonistic effects (Fig. 5c).

The effect of the top/intermediate position is similarly small (χ^2^(15, N=66040)=777.4, *p*<0.001, Cramer’s V = 0.06) and, as for the EXT index, significant differences primarily arise from the predator-prey relationship that may develop, especially when at least one of the invaders acts as a top predator in the separate invasion scenario (Fig. 5d): the frequency of the antagonistic effect is increased by the presence of a predator-prey link.

The impact of trophic position on additivity is further supported by the invaders’ TL index. In line with the results showed in the previous paragraphs (Fig. 5), invaders entering at lower trophic levels are more likely to cause extinctions within the resident food web. Conversely, at higher trophic levels, the resident community exhibits greater resilience, increasing the probability of remaining intact. However, most invasions result in a net reduction in the biomass of the resident food web, when invaders establish at intermediate trophic levels, the probability of a net biomass increase rises (see section S4. of SM1).

When we investigate the variation in frequencies of each additivity category against the TLs of the two invaders (expressed as a percent change relative to the frequencies in the total sample, see Fig. 4), the differences become more visible (Fig. 6). For the EXT index (Fig. 6a), the pattern reveals again, that: i) when both invaders are at higher TL, the resident food web is more likely to remain intact; ii) the likelihood of non-additive outcomes increases as the invaders appear at lower trophic levels, and iii) the frequency of the additive outcomes increases if the difference between the invaders’ TL is higher. In contrast, for the BM index (Fig. 6b), the visualized patterns indicate that invaders at higher trophic levels are associated with an increased probability of additive outcomes. Similarly, the likelihood of synergistic cases also rises with higher trophic levels. The reverse outcome is rare overall but occurs with the highest probability when invaders appear at intermediate trophic levels. This finding aligns with the observation that the highest frequency of “reversed” cases occurs when both invaders are omnivores (Fig. 5c). The most frequent outcome in this case is antagonistic, and the presence of invaders at lower trophic levels further amplifies this tendency. The Welch’s ANOVA test results, examining the influence of TL differences and averages on additivity, are detailed in section S4. of SM1. and Tables S3. in SM2.

**Figure 6.**
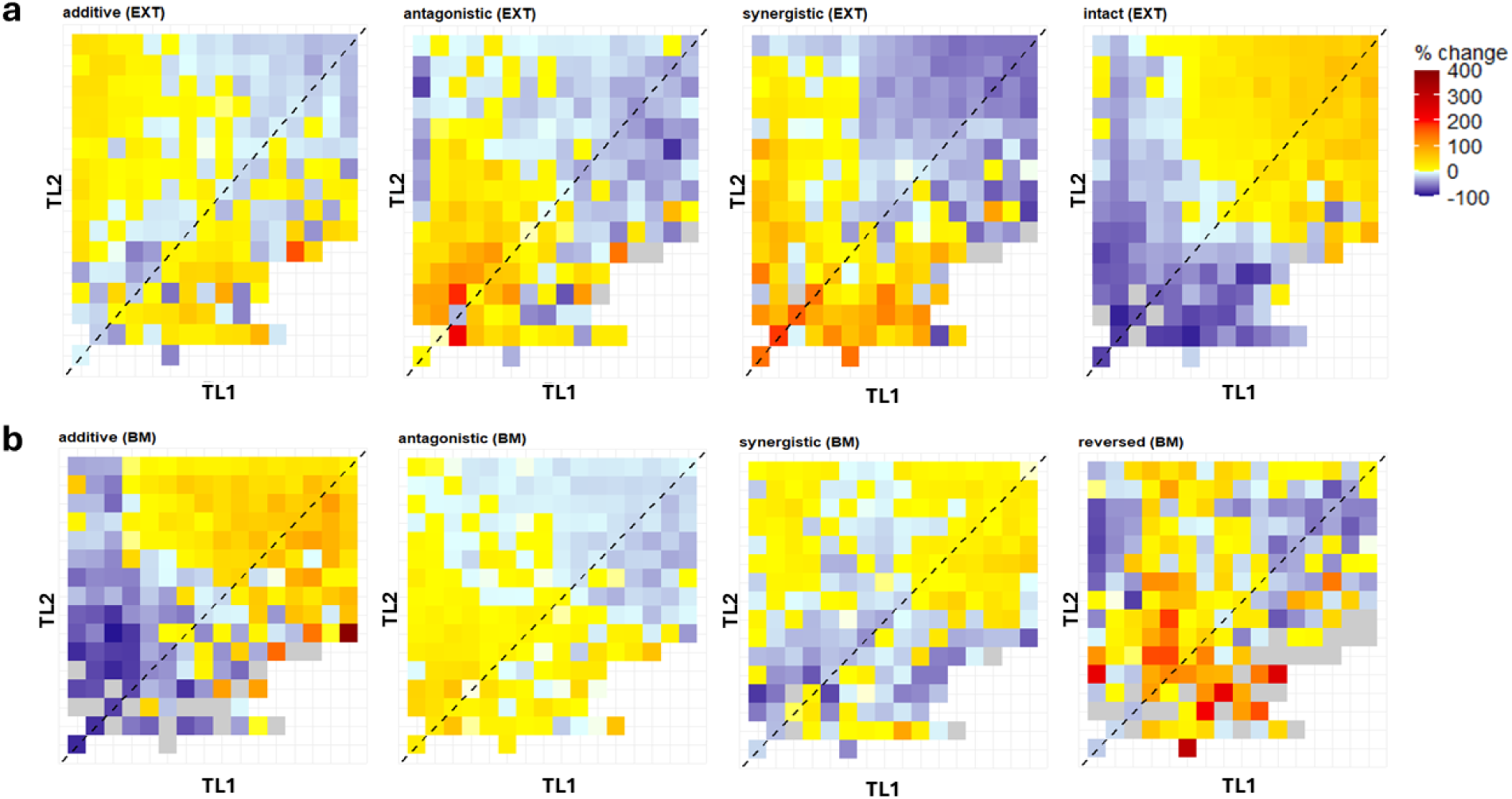
Relative changes of the frequencies of the additivity-categories compared to the overall frequencies (Fig.4a and 4b) in function of the invaders’ trophic level (TL1 and TL2) in case of a) EXT index and b) BM index. Dashed lines represent cases where the invaders’ trophic levels are nearly the same. Index values are normalized between 0 and 1 and grouped in bins of 0.05. Grey dots indicate the absence of a specific category for that TL combination.

### 3.3 Impact of the direct and indirect relationships between invaders

The effect of TL differences and averages on the frequency of each outcome is not independent of the direct/indirect ecological relationship that presents between the two invaders during co-invasion. The invaders’ TL and their TL differences play a significant role in determining the type of ecological relationships between them: a simple predator-prey relationship or trophic cascade typically implies larger TL difference, whereas competitive relationship, particularly for shared prey, is more common when the TL difference is smaller. (section S5. of SM1).

The topological distance between the two invaders has a small but statistically significant effect on the frequency of the different additivity categories for both indices (EXT: χ^2^(6) = 1207.6, *p*<0.001, Cramer’s V = 0.1; BM: χ^2^ (6) = 864.3, *p*<0.001, Cramer’s V = 0.08). Despite the small effect sizes, some clear trends can be observed for both indices (Fig. 7a,d): the establishment of a predator-prey (1 step) relationship between the invaders significantly increases the likelihood of an antagonistic outcome compared to its overall frequency (Fig 4.). However, as the topological distance between the invaders grows, the frequency of synergistic joint effects also increases. Investigating the direct relationship-categories, the frequency of antagonistic outcomes increases further for both indices when the relationship between the invaders is not merely a *predator-prey* (PP) but an *intraguild predator* (IGPred) connection, meaning that the invaders share a common prey (Fig.7b,e). This effect is moderate in magnitude (EXT: Cohen’s ω=0.36; BM: Cohen’s ω = 0.31).

**Figure 7.**
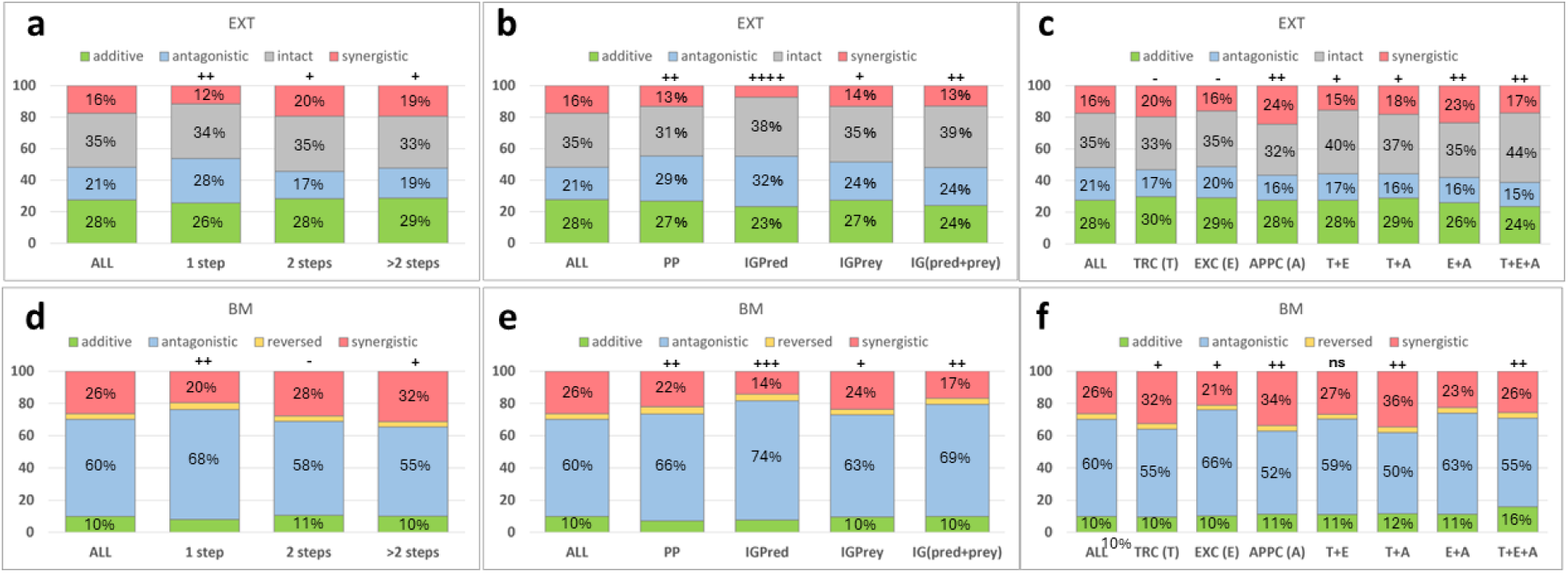
Frequency distributions of the different additivity categories in function of the invaders’ topological distance and the different types of the ecological relationships between them in case of EXT index (a-c) and BM index (d-f). The first columns on the plots show the frequency distribution in the whole dataset (ALL), while the symbols above the rest of the columns indicate whether the effect size (Cohen’s ω) is negligible (-), small (+), moderate (++) or medium (+++) or large (++++), indicating the magnitude of the deviation from the full sample. ns means that the difference is not statistically significant. Frequency-values below 10 % are not displayed on the bars.

For two-step indirect relationships (Fig. 7c,f), the χ^2^-test also shows a weak association between the frequency of additivity categories and different relationship types (EXT: χ^2^(18) = 321, *p*<0.001, Cramer’s V = 0.06; BM: χ^2^(18) = 601.5, *p*<0.001, Cramer’s V = 0.08), indicating that for both indices, exploitative competition (*ExC*) between the invaders increases the frequency of antagonistic and apparent competition (*AppC*) increases the frequency of synergistic outcomes. The trophic cascade (*TrC*) relationship also contributes, albeit to a lesser extent, to an increase in the frequency of synergistic outcomes. (Fig. 7c,f). Detailed results of the χ^2^ tests and the post hoc z tests can be found in Tables S2 in SM2.

### 3.3 Combining the effect of trophic level and the invaders’ direct/indirect relationships

When trophic levels are combined with the topological distance and the direct/indirect connections between the invaders, the picture becomes more nuanced. In Fig. 8., we present the main types of relationships (*PP*, *TrC*, *ExC*, *AppC* and >2 steps connection), while the effects of combinations of these (e.g. *IGPred* or *IGPrey*, etc.) are detailed in the figures in section S6. of SM1. One-step predator-prey relationships consistently increase the frequency of antagonistic outcomes, but this effect is more pronounced when the invaders enter at lower trophic levels, particularly for the EXT index, where the relative increase is more significant. This trend is true for the BM index, too; however, there is a higher baseline probability of antagonistic outcomes, resulting in a smaller percent change (Fig. 8a). Measuring species extinctions (EXT index), if the *PP* relationship is combined with exploitative competition, this effect is even stronger (*IGPred*). However, if they share a common predator (*IGPrey*), the probability of antagonistic outcomes tends to decrease at higher trophic levels (Fig S5a). For the BM index, these two combined categories lead to a slight decrease in the frequency of antagonistic outcomes at higher trophic levels, where the frequency of additive outcomes increases significantly (Fig. S6a).

**Figure 8.**
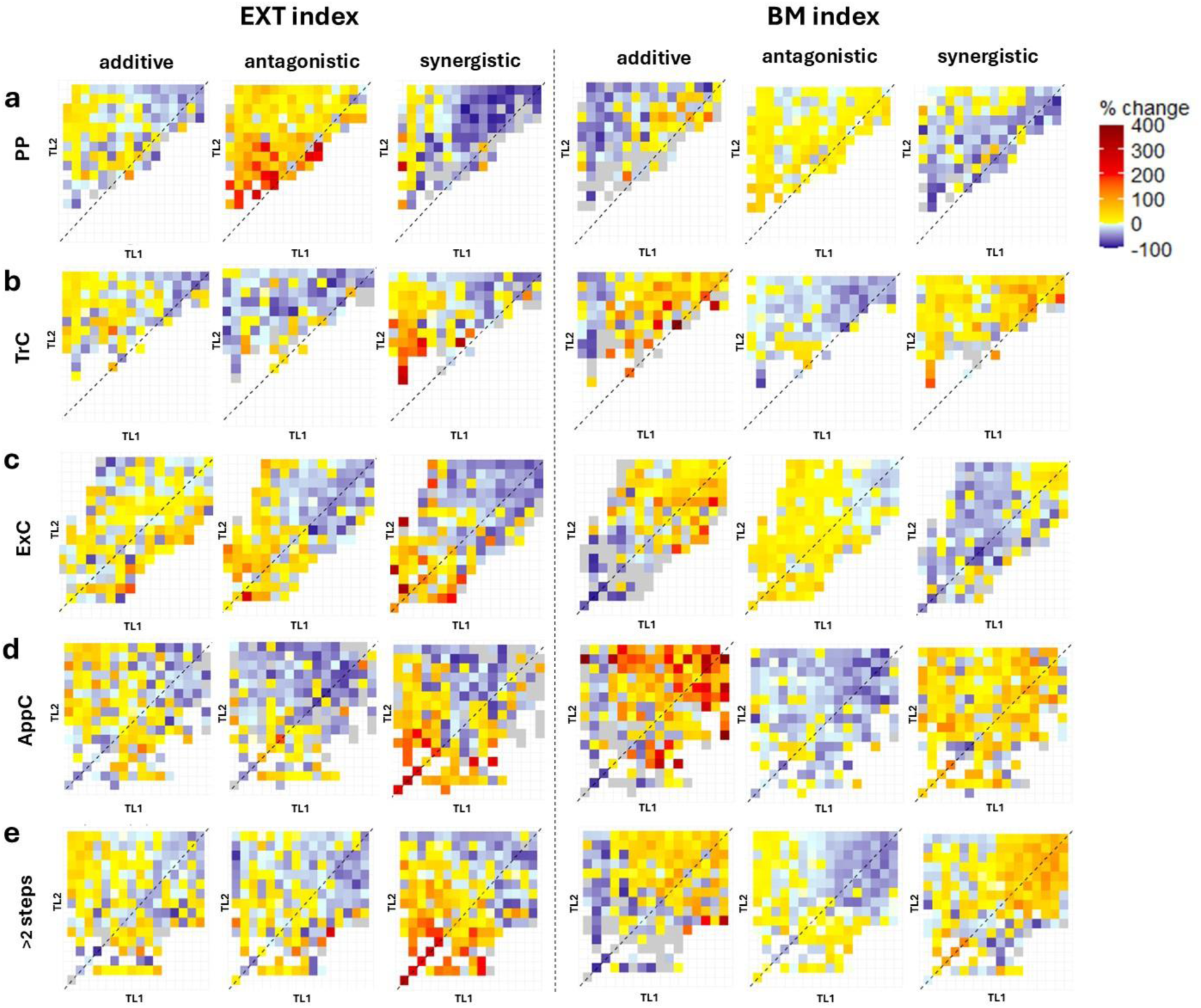
Relative changes of the frequencies of the additivity-categories compared to the overall frequencies (Fig.4a and 4b) in function of the invaders’ trophic level (TL1 and TL2) and considering the main direct and indirect connection-categories between the invaders. The dashed lines indicate the cases when the invaders’ trophic levels are (almost) the same. Dashed lines represent cases where the invaders’ trophic levels are nearly the same. Index values are normalized between 0 and 1 and grouped in bins of 0.05. Grey dots indicate the absence of a specific category for that TL combination.

Indirect two-step relationships increase the frequency of additive outcomes in the most cases for case of both approaches, but for the BM index this increase is only observed at higher trophic levels, while at lower trophic levels the frequency of additive outcomes decreases or even disappears (Fig. 8b-d). For the EXT index, two-step indirect relationships show an increase in the frequency of non-additive outcomes at lower trophic levels, which is more pronounced in the case of synergism. This trend is particularly notable for the *TrC* and *Appc* categories, where the probability of antagonistic outcomes decreases while that of synergistic outcomes increases significantly at the lowest trophic levels. For the BM index, the effects of the two types of indirect competition diverge more significantly. At higher trophic levels, additive effects become more frequent for both types of competition, especially for the *AppC* relationship. Meanwhile, *ExC* increases the probability of antagonistic outcomes, whereas *AppC* increases the probability of synergistic outcomes. Similar to the EXT index, the *TrC* relationship increases the frequency of both additive and synergistic outcomes; however, the increase in synergistic outcomes occurs not only at lower trophic levels but across the whole TL spectrum.

When the two invaders are even more distantly connected within the community (>2 steps), the probability of antagonistic outcomes increases with larger trophic level differences, especially for the BM index (Fig 8e). Synergistic cases show a different pattern for the two metrics. For the EXT index, their frequency rises with increasing topological distance, with the most significant increases observed at lower and close/same trophic levels. Interestingly, synergism is the most frequent when the two invaders occupy the same, low trophic level within the food web, but are separated by a large topological distance. Conversely, in case of the BM index, the frequency of synergistic outcomes increases more prominently at higher and more similar trophic levels.

## 4. DISCUSSION

With accelerating invasion rates across ecosystems, interactions between invasive species are becoming more prevalent, underscoring the urgency of understanding the effects of their direct and indirect relationships (Rauschert and Shea, 2017). These interactions not only influence the success of the invasion (Jackson, 2015; Johnson et al., 2009; Móréh et al., 2024) but also significantly impact the resident community. However, the complexity and diversity of these interactions make it challenging to predict the direction and the magnitude of the potential impact of two or more invaders.

Most studies on invaders have focused on their impacts at the level of individuals or populations of particular native taxa, often neglecting their broader effects on entire communities expressed by, for example, taxonomic richness and various indices of diversity (Parker et al., 1999). We investigated the joint impact of two invaders on whole food webs using two metrics: the change in the diversity of the resident community, measured by the number of species extinctions, and the net biomass changes, which were recorded and compared across the two invasion scenarios (separated and simultaneous). In the extinction-focused analysis, approximately one-third of the outcomes resulted in an intact network (no extinctions in either scenario). However, the frequencies of the remaining three outcomes show only slight differences. The relatively higher proportion (28%) of additive outcomes can be attributed to scenarios where only one of the two invaders causes extinction, making it less likely for their combined effect to deviate from the sum of their individual effects. When focusing on biomass change, antagonistic outcomes are the most prevalent (60%), synergistic outcomes occur less frequently (26%), and additive outcomes are comparatively rare (10%). In this approach, it is also possible for the joint effect of the two invaders to have an opposite sign to their individual effects; however, this was a relatively rare occurrence in our study (4%).

Our results demonstrate that the choice of metric used to describe invasion effects significantly influences the observed nature of the joint effect, as the frequency of different outcomes varies notably between the two approaches. Here, this discrepancy may arise because successful invaders always cause some degree of biomass change, whereas diversity may remain unaffected, and extinction does not always occur. While our findings are consistent with previous studies showing that additive and antagonistic outcomes are more common than synergistic ones (Jackson, 2015; Jackson et al., 2016), meta-analyses have also highlighted that outcomes in individual case studies can deviate considerably from global trends (Jackson, 2015), an observation that is again consistent with our findings. The common effects may differ depending on whether the focus is on species-level (Braga et al., 2020; Nystrom et al., 2001) or community-level traits (Johnson et al., 2009; Levin et al., 2002). This is one of the reasons why predicting the outcome of a multispecies invasion is challenging (Crain et al., 2008). On the other hand, field and mesocosm experiments (Preston et al., 2012) and meta-analyses (Cameron et al., 2016; Thomsen et al., 2014) have demonstrated that the effects of invaders are significantly influenced by their trophic positions. Thomsen et al., (2014) found that marine invaders negatively affected biodiversity within similar trophic levels through competition and consumption, but their impacts were less negative or even positive at higher trophic levels where they served as a food source (*relative trophic position hypothesis*). We found that the lower the trophic position (or TL index) at which the invader enters the system, the more likely it is to cause extinctions in the resident network, or conversely, entering at a higher trophic level, the network is more likely to remain intact in this respect. Similarly, the biomass-reducing effect is also most pronounced at low trophic levels, but we found that entering at intermediate trophic levels may also cause the increase of total resident biomass. Examining the joint effect in terms of extinctions, we find that the lower the trophic positions of the two invaders and the smaller the difference in their trophic levels, the higher the probability of non-additive outcomes. Conversely, a larger trophic distance between invaders increases the likelihood of additive outcomes. If invasions do happen at higher trophic positions, the network is more likely to remain intact. On the other hand, when considering biomass change, the frequency of antagonistic outcomes, which are already prevalent, further increases if the invaders enter at lower trophic levels, whereas their higher trophic levels tend to favor additive or synergistic outcomes.

In their meta-analysis, Cameron et al., (2016) observed that the trophic position of terrestrial invertebrate invaders significantly influenced their impact on species abundance, with predators exerting stronger negative effects than omnivores or herbivores. Consistently with this finding, we demonstrated that an invader exerts a stronger impact when it acts as a top consumer compared to when it occupies an intermediate position, i.e., when it has a predator (either a resident or another invader species). A top invader causes more species extinctions on average than intermediate invaders, and it also induces greater changes in biomass in both negative and positive directions. Nevertheless, the fact that invaders are intermediate or top consumers does not, by itself, significantly affect the likelihood of different additivity categories, regardless of the response metric considered (EXT or BM indices). However, if one invader preys on the other invaders, the probability of an antagonistic joint effect increases significantly. Interestingly, this effect is significant only when at least one of the two invaders acts as a top predator, whereas if both are in intermediate positions, the effect is minimal. This highlights the critical importance of considering the direct or indirect relationships between invaders when examining their joint effects.

Several field and mesocosm experiments have been conducted to investigate the impact of direct and indirect ecological relationships between invaders. As mentioned, invaders can negatively affect one another through predation, thereby reducing or minimizing their combined impacts on native biota (Ross et al., 2004). In a study on interspecific predation between two invasive crustacean species, Buřič et al., (2009) also demonstrated that both invaders could negatively affect each other through intra-guild predation. Similarly, the investigations of co-occurring crayfish invaders revealed that competitive interactions between them may have long-term negative effects on at least one of the species (Hudina et al., 2011). When invasive species establish themselves at multiple trophic levels, they can create a trophic cascade that may facilitate the success of the invaders, leading to synergistic joint effects (Crego et al., 2016; Liversage et al., 2021). These examples underscore the critical need to comprehend interspecies interactions among invasive species. Examining the direct and indirect relationships between invaders using both extinction and biomass metrics, we also showed these basic correlations, namely that predator-prey relationship between them increases the likelihood of an antagonistic effect, especially when coupled with competition for a shared resource (intraguild predation). This aligns with the observation that exploitative competition also amplifies antagonistic effects, even in the absence of a direct feeding relationship. Conversely, apparent competition, such as that observed in a trophic cascade, tends to increase the likelihood of synergism.

Given the strong correlation between the difference or similarity in trophic levels and the direct/indirect relationships between the two invaders, examining these factors together provides a more accurate understanding of the probabilities of various outcomes. Trophic level not only determines whether an invader causes extinctions within the food web or decrease/increase its’ biomass but also shapes the extent and direction of how interactions between the two invaders influence the frequency of additive or non-additive outcomes. For example, when considering biomass changes in the resident food web, even if exploitative competition develops between the two invaders, additive outcomes are more likely than antagonistic ones at higher trophic levels. In the case of the extinction approach (EXT index), even if invaders form a trophic cascade, their effect is more likely to be synergistic only if both enter the network at a lower position. On the other hand, the frequency of synergistic outcomes increases most significantly when the two invaders enter the community at the same low trophic level, but a distant, more than two-step relationship develops between them. This strong effect of trophic level is not totally surprising, since it significantly influences invasion success, too (Dijoux et al., 2024; Móréh et al., 2024; Romanuk et al., 2009). It is also strongly correlated with the likelihood of an invader causing extinction: due to the bottom-up effect invaders at lower trophic levels tend to cause more damage to the network. However, the direction of biomass change is less straightforward: we have shown that in addition to the frequent biomass-reducing effect, invaders at intermediate trophic levels can also lead to increases in total biomass. This also means that predicting the overall impact of omnivores invading at intermediate trophic levels on resident communities is challenging, as they can reduce diversity at lower trophic levels through predation and competition, while simultaneously providing resources for higher trophic level predators (Parkos et al., 2003). This is further corroborated by the notable increase in the frequency of the otherwise rare “reversed” outcome at this trophic position, where the direction of biomass change resulting from the joint invasion was opposite to that observed when the invaders acted independently.

Interactions between co-occurring invaders are crucial for wildlife management, however efforts generally focus on monitoring and controlling just one invader at a time (Glen et al., 2013). For example, managing just one invader can significantly accelerate recovery, as reducing one of them in this case of their synergistic common effect may drastically lessen the overall impact on the community (Fong et al., 2018). Conversely, the removal of one invader might unexpectedly harm native species more than when both invasive species are present. Such counterintuitive outcomes likely stem from the positioning and hierarchical relationships of the co-occurring invaders within the food web (Ballari et al., 2016).

As in our previous study (Móréh et al., 2024), we utilized an allometry-based model that, while moderately complex, offers a more accurate representation of species interactions compared to the commonly used Lotka-Volterra models. This approach is particularly suited for aquatic systems, where trophic interactions can often be inferred from allometric relationships between predator and prey body sizes (Vagnon et al., 2022; Woodward et al., 2005). This method has limited applicability to terrestrial ecosystems, where body mass is not always a reliable predictor of plant-consumer interactions. However, efforts are underway to adapt and extend this framework to these communities (Valdovinos et al., 2023). The size of an individual is a key factor in determining its trophic position. Consequently, the ecological roles of the invaders within a community change with body size during their ontogeny, leading to a size-dependent variable mix of competition and predation in trophic interactions (Schröder et al., 2009). This aspect, however, was not accounted for in our current study. Another limitations of our study is that we prioritized the properties and interactions of the two invading species, while neglecting the role that the resident network’s structure can play in terms of the response to the invasion (see Lurgi et al., 2014; Romanuk et al., 2009). Excluding the resident network structure from the analysis enhances our ability to study how invaders potentially most important factors, the trophic positions and ecological interactions modify their effect on the resident food web. Nonetheless, a wider range of species traits and network indices remain to be explored (Gouveia et al., 2021; Móréh et al., 2021; Romanuk et al., 2009), as well as several other metrics for community responses might be studied in the future (Móréh and Jordán, 2019). Acknowledging that the impact of multiple invasions hinges on the balance of relationships between invaders and their interactions with the resident community (Preston et al., 2012), our study advances a theoretical framework for understanding the varied outcomes of single- and two-species invasions within food webs. We emphasize the pivotal role of the invaders’ trophic position and their interspecies interactions, knowing their importance in formulating invasive species management strategies and preserving native and semi-native ecosystems.

## 5. ACKNOWLEDGEMENTS

This work was supported by the European Union’s Horizon 2020 research and innovation programme under grant agreement No 952914. English grammar and style are checked by DeepL Write beta 2.0.

