## Supplementary_Material_1 for "Invaders’ trophic position and their direct and indirect relationship influence on resident food webs"

#### S1. Allometric bioenergetic model (ABM)

The allometric bioenergetic model (ABM) is an extensively utilized framework for modelling various types of food webs, especially aquatic communities. Since we employed the model and its parameters without alterations, this summary is directly adapted from the description provided by Brose et al. (2006).

In the allometric coefficients-updated consumer-source model (Brown et al., 2004), changes in the biomass density ($B_i$) of the species are described by a system of differential equations:

$$\frac{dB_{i(basal)}}{dt} = r_i(M_i)B_iG_i - \sum_{c=cons} \frac{x_c(M_c)y_c}{e_{ci}} B_cF_{ci}$$

$$\frac{dB_{i(consumer)}}{dt} = \sum_{\rho=res} x_i(M_i)\mathrm{B}_iF_{i\rho} - \sum_{c=cons} \frac{x_c(M_c)y_c}{e_{ci}} B_cF_{ci} - x_iB_i$$

The first equation refers to the basal species, where the first term on the right-hand side describes the basal species' growth. The mass-specific maximum growth rate of species *i* is represented by $r_i(M_i)$, and $G_i$ is the term for the logistic net growth rate ($G_i$ = 1 - ($B_i$ / $K_i$), where $K_i$ denotes the carrying capacity associated to species *i*). The second term expresses the biomass-decrease of the basals due to consumption: $x_c(M_c)$ represents the consumers' mass-specific metabolic rate, and $y_c$ denotes the maximum consumption rate relative to its metabolic rate. Finally, $e_{ci}$ represents species *c* assimilation efficiency when consuming species *i*. In the second equation, which applies only to consumptive species, the increase in biomass comes only from consumption (first term on the right hand side), but the decrease comes not only from consumption by predators (second term), but from a mortality of its own, proportional to the mass-dependent metabolic rate (last term).

The nonlinear functional response terms represent the actual consumption rate realized when species *i* preys upon species *j*:

$$F_{ij} = \frac{B_j^h \omega_{ij}}{B_0^h + \sum_{k=res} \omega_{ik} B_k^h}$$

where $\omega_{ij}$ represents the relative consumption rate of species *i* when consuming species *j*, $B_0$ is the half saturation density, and *h* is the Hill-coefficient. For consumers with *n* resources, we use uniform relative consumption rates ($\omega_i = 1/n_i$). We set $h = 1.5$, which compromise the type II response ($h = 1$) and type III functional response ($h = 2$).

The central concept of the model is based on the negative quarter-power scaling relationships that link biological rates of production (*R*), metabolism (*X*), and maximum consumption (*Y*) to the body mass of the species:

$$R_P = a_r M_P^{-1/4}, X_C = a_x M_C^{-1/4}, Y_C = a_x M_C^{-1/4}$$

where $a_r$, $a_x$ and $a_y$ are allometric constants, *C* and *P* indicate consumers' and primary producers' parameters, respectively (Brown et al., 2004; Yodzis and Innes, 1992). By setting the growth rate of the basal species to unity ($r_{basals} = 1$), we establish the time scale for the system. The metabolic rates of the species are then normalized relative to this time scale:

$$x_i = \frac{X_C}{R_P} = \frac{a_x}{a_r}\left(\frac{M_C}{M_P}\right)^{-1/4}$$

while the maximum consumption rates are normalized by the metabolic rates:

$$y_i = \frac{Y_C}{X_C} = \frac{a_y}{a_x}$$

We leverage the observation that, particularly in marine food webs, predators are typically larger than their prey (Ou et al., 2017). The predator-to-prey body-mass ratio is a key metric for analyzing trophic structure and community dynamics (Trebilco et al., 2013; Perkins et al., 2022; Vagnon et al., 2022). We define a constant predator-to-prey body mass ratio (Z), where a value of $Z = 10^2$ indicates that the predator is 100 times heavier than its prey (Brose et al., 2006). As a result, we establish a relationship between a species' body mass and its trophic level (TL), expressed as $M_C = Z^{TL}$.

Within this framework, the body masses of all predators are defined relative to the body mass of the basal species. This method accounts for variations in body mass across trophic levels and provides valuable insights into the size structure and dynamics of predator-prey interactions within the food web.

| ***Parameter*** | ***Description*** | ***Fixed value*** |
|---|---|---|
| $\boldsymbol{B_i}$ | biomass of *i* | $B_{initial} = 1$ |
| $\boldsymbol{r_i}$ | mass-specific growth rate | 1 |
| $\boldsymbol{K_i}$ | carrying capacity | 1 |
| ***Z*** | consumer-resource body mass ratio (BMR) | 100 |
| $\boldsymbol{x_i}$ | mass-specific metabolic rate | $\frac{a_x}{a_r}\left(\frac{M_C}{M_P}\right)^{-1/4}$ |
| $\boldsymbol{a_r}$ | allometric constant | 1 |

| | | |
|---|---|---|
| $a_x$ | allometric constant | 0.6 |
| $y_i$ | max. consumption rate | 6 |
| $e_{ci}$ | assimilation efficiency of *c* when consuming *i* | 0.65 |
| $B_0$ | half-saturation biomass | 0.5 |
| $\omega$ | relative consumption rate | $1/n_i$ |
| $h$ | Hill-coefficient | 1.5 |

*Table S1. Model parameters; C: consumers; P: primary producers; $n_i$: number of i's resources; $a_x$, $y_i$ and $e_{ci}$ represent the mean of the constants given for vertebrate/invertebrate and herbivorous/carnivorous species*

**S2. Simulation of invasion**

The invasion was simulated exactly as described in our previous work (Móréh et al., 2024). We extended the previously constructed resident webs to include the invading species, depending on the specific scenario under investigation (either separate or joint invasion). The links for the invaders were randomly assigned according to the same criteria used to generate the resident food web. Our question is how the outcome of a joint invasion differs from when the same invaders are introduced separately into the system. For practical reasons, we reversed the typical process of expanding the generated food webs. First, we simulated the joint invasion, and in the subsequent round, we re-ran the simulation with the same invaders, but their previously assigned links were reintroduced separately into the resident networks. This reverse ordering was essential to capture any potential predator-prey relationships between the two invaders during the analysis. By allowing these relationships to emerge randomly during the link assignment in the joint invasion simulation, we avoided artificially creating them afterward. This approach enabled us to naturally investigate the distinct impacts of the invaders while accounting for their potential interactions. Although the invaders appeared almost simultaneously in the simulation with a very short time lag, for practical reasons they were labelled I1 and I2 to indicate the order of their appearance. The pairing process involved placing I1 directly above the basal species ($I1_{\mathrm{Nbasal+1}}$), fixing its position, and then placing I2 next to it at each subsequent level ($I2_{\mathrm{I1+1...S+2}}$). For example, if a resident network contained S species with three basal species, I1 would be the first invader, positioned as species #4, and would be systematically paired with I2, which would be placed at position #5, and so on until #S+2. After each "round", I1 would move up one position and be paired with I2, positioned directly above it, until both invaders reached positions #S+1 and #S+2. This technical solution ensures that invader pairs are systematically distributed across the entire consumer spectrum within the network. For each pair of invaders, we have drawn randomly 10 different link-sets. If the resulting combination led to a stable state by the end of the simulation, the combination was kept; otherwise, it was discarded, and a new set of links was drawn. More complex stable states, such as limit cycles or chaos, were not observed. This process generated a total of 121,000 invasion scenarios that could be tested in 110 resident food webs. Since we focused in this study on how the combined effect of invaders on the resident network differs from the effect of invaders separately, we only considered invasion results where both successfully invaded in both scenarios, i.e., established a stable population in the network. This represents an analysis of the impact of 66040 invasion events.

### S3. Statistical analyses

We applied $\chi^2$ tests to examine the association between the feeding types of the two invaders, their position in the network (either top or intermediate), and the additivity categories. As expected with the large sample size, the results were statistically significant (Cohen, 1988). To identify the specific differences driving this significance, a Bonferroni-corrected post-hoc z-test was performed, with Cramer's V used to measure the strength of the associations. Additionally, a proportion test (goodness of fit) was conducted to determine whether the frequencies of the different effect types deviated significantly from the overall probabilities observed in the full dataset, with Cohen's w used to express effect size. The same methodology was employed to investigate whether the direct/indirect ecological relationships between the two invaders influenced the type of combined invasion effect observed.

We analyzed the relationship between the invaders' trophic level (TL) indices by examining both their difference and mean, recognizing that these two metrics are interrelated. While small differences between TL indices can correspond to both low and high averages, larger differences tend to align the average toward a central value. Consequently, we analyzed these metrics separately rather than combining them into a single measure, using their absolute values to quantify differences. To evaluate how the differences or means of the indices varied across different effect types, we applied a Welch ANOVA test, chosen due to the unequal variances in the samples. Square root transformations were performed on the variables to meet normality assumptions. Effect sizes ($\eta^2$ and Cohen's *d*) were used to assess the strength of relationships. For detailed comparisons between groups, we employed a Games-Howell post hoc test, which is well-suited for multiple category comparisons.

A comprehensive summary of the statistical results is provided in the tables of the Supplementary Material 2.

### S4. Impact of the invaders' trophic position

Both the success of the invasion itself and its impact on the resident food web are strongly influenced by the trophic position of the invader (Romanuk et al., 2009; Thomsen et al., 2014). Looking at the number of extinctions, we find that the lower the trophic position of the invader, the more likely it is to cause extinctions in the resident network (Fig. S1a). In terms of biomass variation, we know that in most cases, especially at low trophic positions, the invader causes a stronger decrease in total biomass that is not independent of a higher extinction probability (Fig. S1c). However, the impact differs if the invader itself has a predator (either resident or another invader) compared to top consumer invaders at the same trophic position. Our findings indicate that the effect of an invader acting as a top predator is stronger than that of invaders occupying intermediate trophic positions in both extinction rates and biomass changes (Fig. S1b, d.).

Similarly, the invaders' TL index also significantly influences the likelihood and nature of its impact. In 22% of all cases studied, both invaders independently caused extinctions during separate invasions. In 38% of cases, only one invader led to extinction, while in the remaining 40%, no extinctions occurred (see Fig. 4a and 4b in the main text). As illustrated in Fig. S2, invaders entered at lower trophic levels are more likely to cause extinctions within the resident network. Conversely, at higher trophic levels, the resident network is more resilient, making it more likely to remain intact. On the other hand, most invasions result in a net reduction in the biomass of the resident network. When invaders establish themselves at intermediate trophic levels, there is an increased probability that the invasion will lead to a net increase in the resident network's biomass, as it can provide an additional resource for resident predators at higher positions than the invader. These patterns underscore the importance of an invader's trophic position in determining both its direct and indirect impacts on the ecosystem.

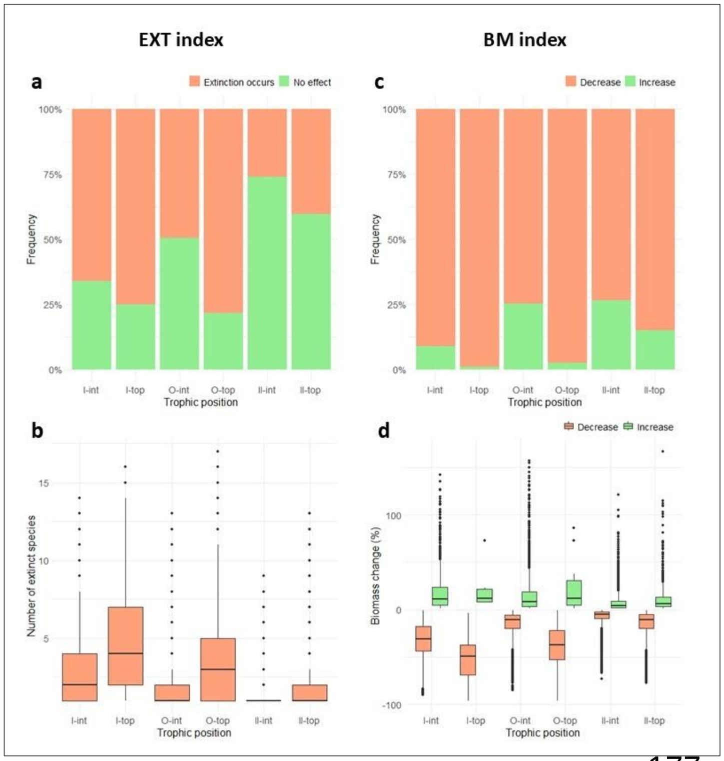


***Figure S1.*** *Effect of invasion on the community based on invaders' trophic position for EXT (a, b) and BM indices (c, d). Bar plots depict the frequency (%) of invasion events causing or not causing extinction (a) and decreasing or increasing total resident biomass (c). Box plots illustrate the effect's magnitude through the distribution of extinct species numbers (b) and biomass change (%) (d).*

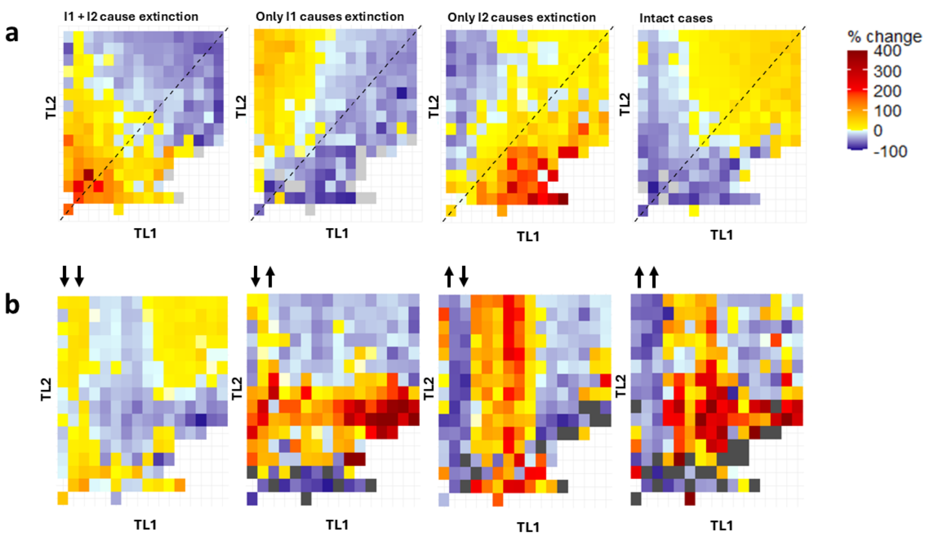


***Figure S2.*** *Relative changes of the frequencies of the effect- and direction-categories compared to the overall frequencies (Fig.4 in the main text) in function of the invaders' trophic level (TL1 and TL2) for a) EXT index and b) BM index, where the arrows indicate the direction of the biomass change. Dashed lines represent cases where the invaders' trophic levels are nearly the same. Index values are normalized between 0 and 1 and grouped in bins of 0.05. Grey dots indicate the absence of a specific category for that TL combination.*

When examining the TLs' difference and average, the Welch ANOVA test indicates that for the EXT index, while there are statistically significant differences between additivity categories from this point of view, the effect sizes ($\eta^2$) vary: for the TL difference, the effect size is small ($F$(3,32206)=149.3, $p$<0.001, $\eta^2$=0.007); for the mean TL, the effect size is medium ($F$(3,31692)=2320, $p$<0.001, $\eta^2$=0.093). The post hoc test reveals that the most pronounced differences occur between the intact and synergistic groups. Specifically, the average TL of the invaders is much lower in synergistic cases compared to situations where the invasion does not result in extinction ($M$=-0.14, $p$<0 .001, Cohens' $d$ = -0.88). These results are illustrated in Fig.S3a.

Similarly, for the BM index (Fig. S3b), while statistically significant results were obtained, the effect sizes are minimal: the TL difference has a negligible effect size ($F$(3,8950) = 30.8, $p$<0.001, $\eta^2$ = 0.001), and the mean TL has a small effect size ($F$(3,9031) = 588.6, $p$<0.001, $\eta^2$ = 0.021). The post hoc test revealed that in this case, the additive outcome group differs significantly from the others: the largest difference is with the antagonistic group ($M$=-0.08, $p$<0.001, Cohen's $d$ = -0.5), a slightly smaller difference is observed with the reversed group ($M$=-0.07, $p$<0.001, Cohen's $d$ = -0.45), and, the smallest difference is with the synergistic group ($M$=-0.05, $p$<0.001, Cohen's $d$ = -0.3). The negative effect sizes indicate that the average TL of the invaders is higher in additive cases compared to the other outcomes.

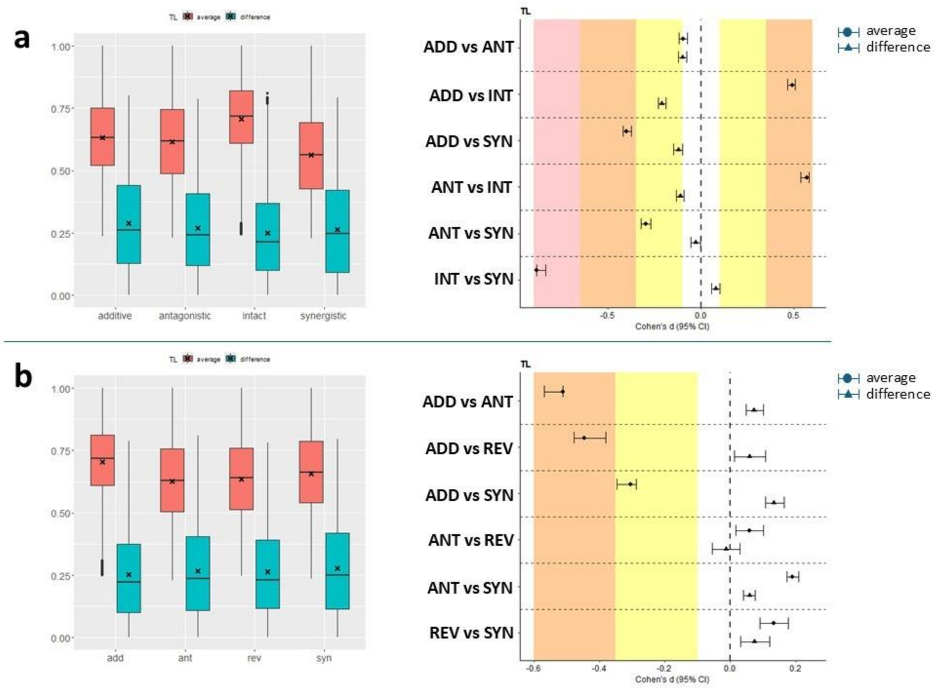


***Figure S3*** *The invaders' TL indices are compared based on their differences (blue) and averages (red) across different additivity categories (left panels). The differences between these categories are represented by Cohen's d effect size metric with 95% confidence intervals (right panels). The background color indicates the magnitude of the difference: negligible (white), small (yellow), medium (orange), or large (red). a) EXT index, b) BM index.*

**S5. Connection between the invaders' trophic level and their direct/indirect relationships**

The trophic levels of the invaders and the differences between them play a key role in shaping the type of ecological relationships that can emerge between them: when both invaders occupy the same or similar trophic levels, they are more likely to engage in competitive interactions, such as exploitative competition (*ExC*, competing for the same prey) or apparent competition (*AppC*, sharing a common predator). Greater differences in trophic levels increase the likelihood of developing hierarchical relationships, such as predator-prey interactions or cascading effects within the network.

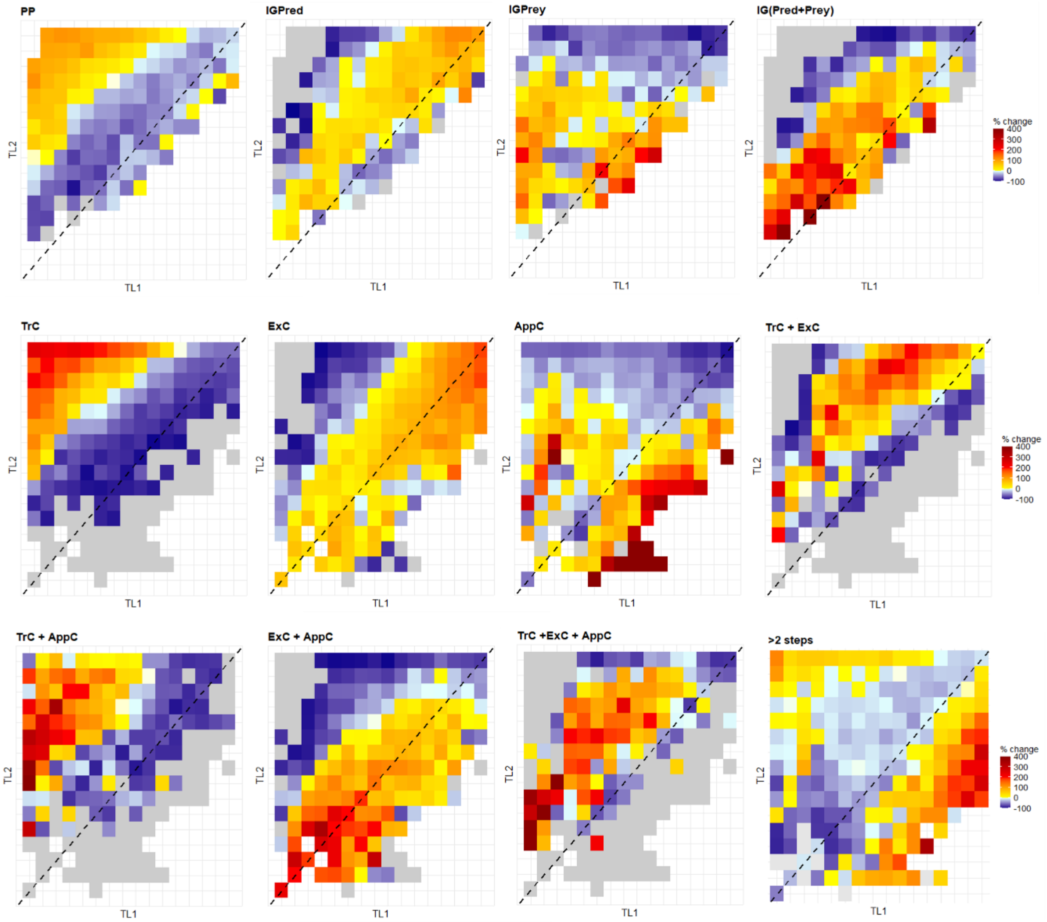


***Figure S4.*** *Relative changes of the frequencies of the direct and indirect relationship-categories compared to their overall frequencies (Fig.3 in the main text) in function of the invaders' trophic level (TL1 and TL2). Dashed lines represent cases where the invaders' trophic levels are nearly the same. Index values are normalized between 0 and 1 and grouped in bins of 0.05. Grey dots indicate the absence of a specific category for that TL combination.*

**S6. Joint effect of the invaders' TL and their direct/indirect ecological relationship on the frequency of additivity categories**

Figures S5. and S6. show the relative changes of the frequencies of the additivity-categories compared to their overall frequencies (see Fig.4 in main text) in function of the invaders' trophic level (TL1 and TL2) *and* their direct and indirect ecological relationship for the EXT (Fig S5.) and BM (Fig. S6) indices, respectively. Dashed lines represent cases where the invaders' trophic levels are nearly the same. Index values are normalized between 0 and 1 and grouped in bins of 0.05. Grey dots indicate the absence of a specific category for that TL combination.

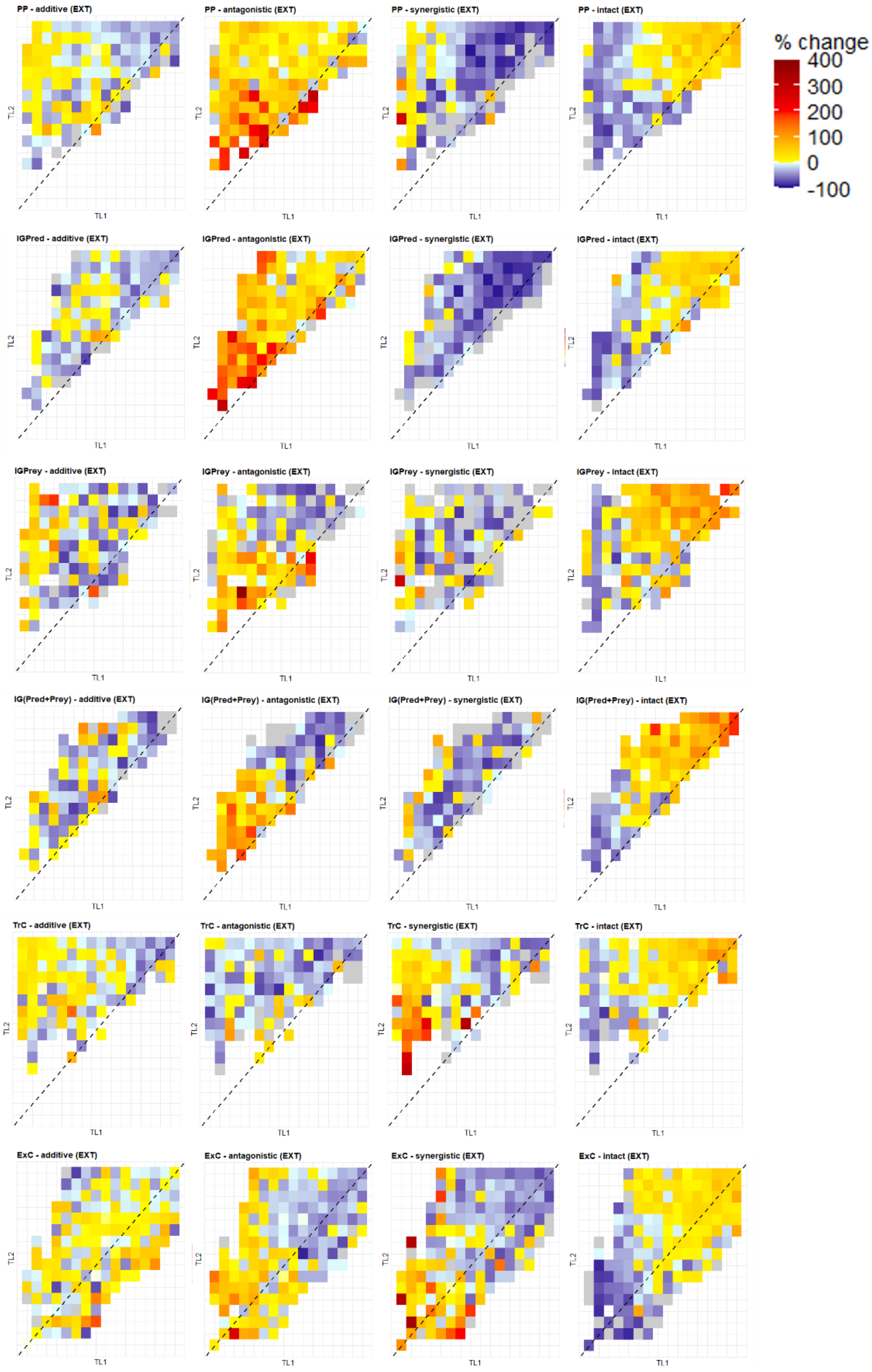


***Figure S5*** *Relative frequency-changes of the different additivity categories vs the invaders' TL indices and their direct/indirect connections in case of EXT index.*

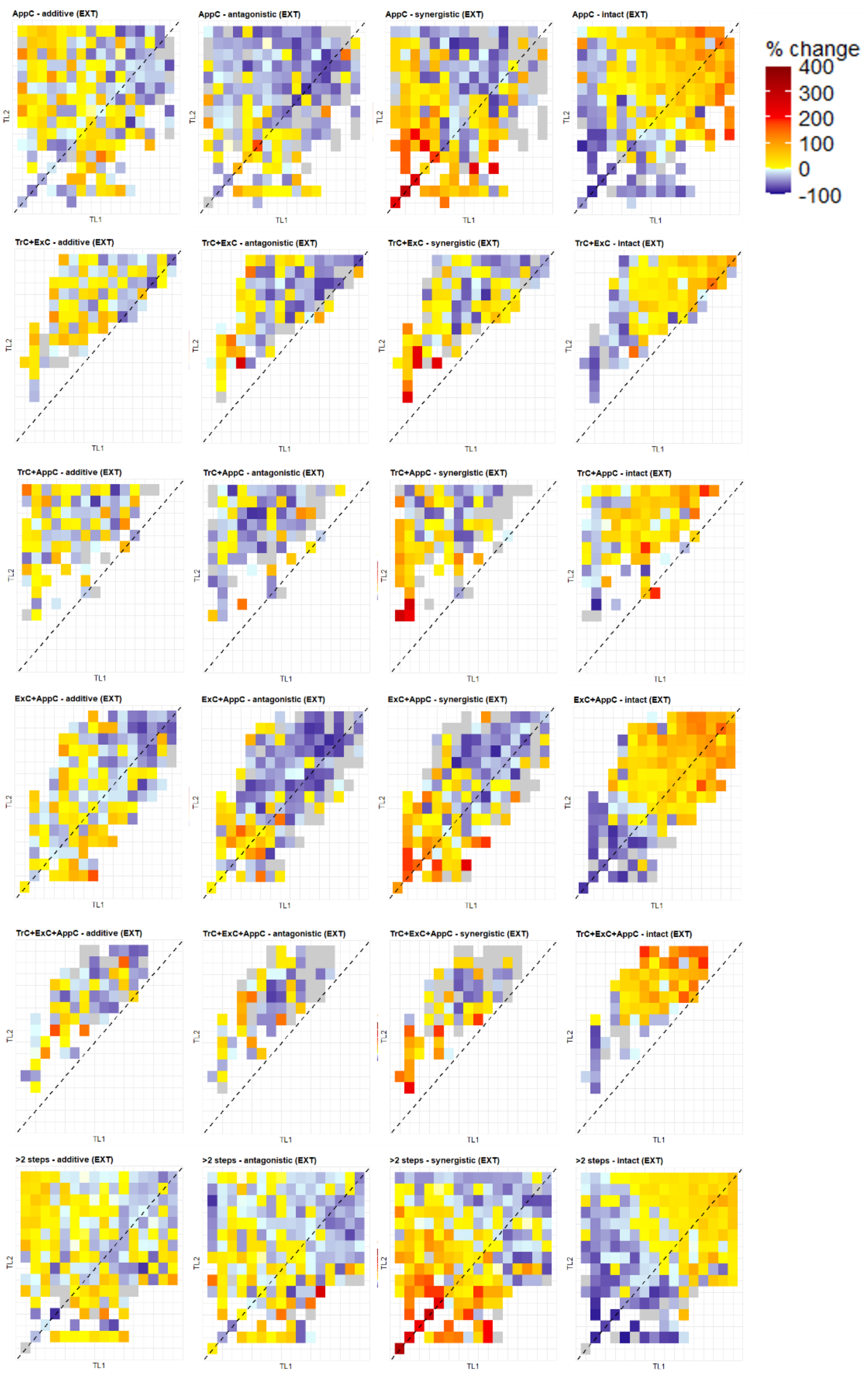


***Figure S5*** *(continued).*

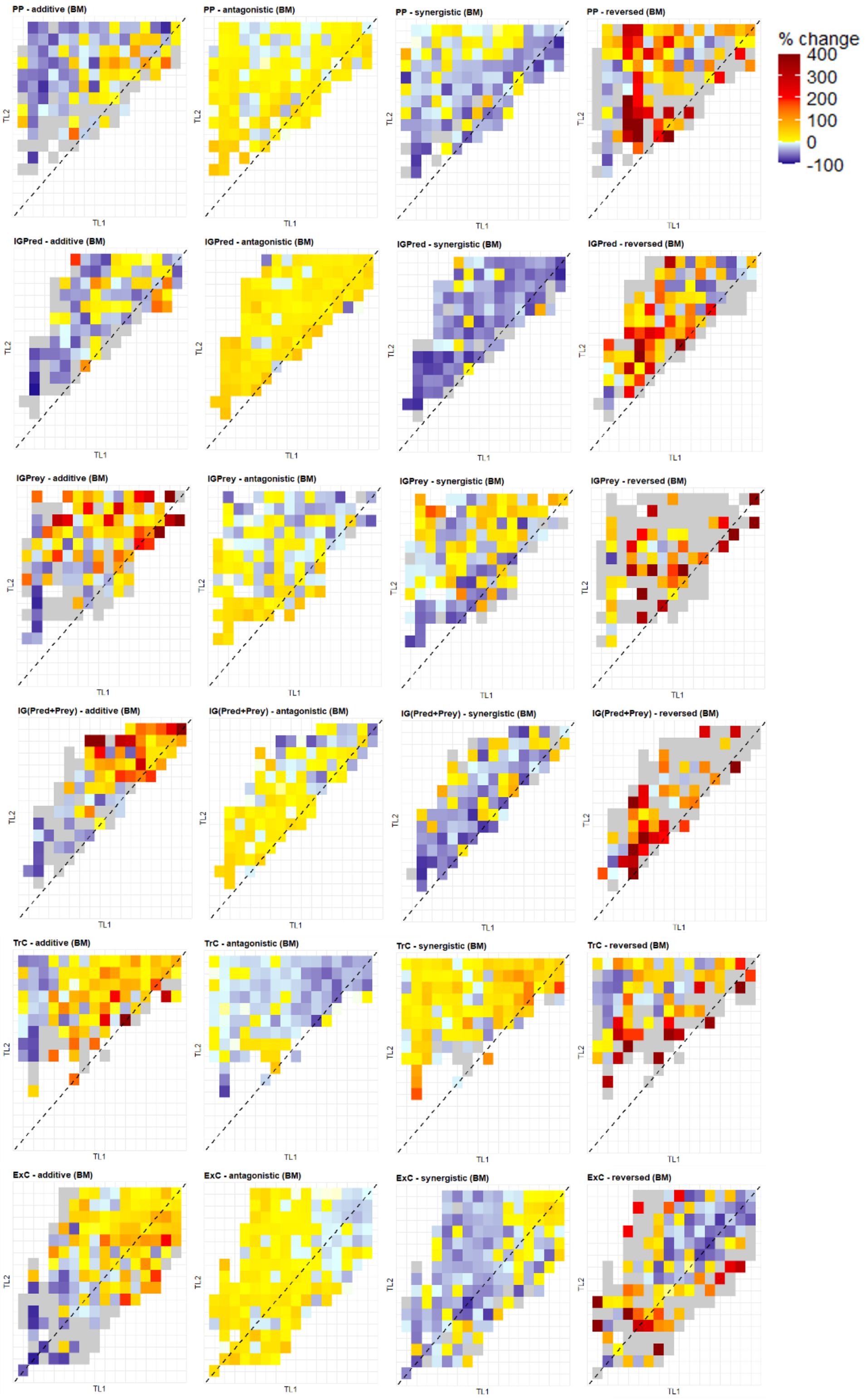


***Figure S6*** *Relative frequency-changes of the different additivity categories vs the invaders' TL indices and their direct/indirect connections in case of BM index.*

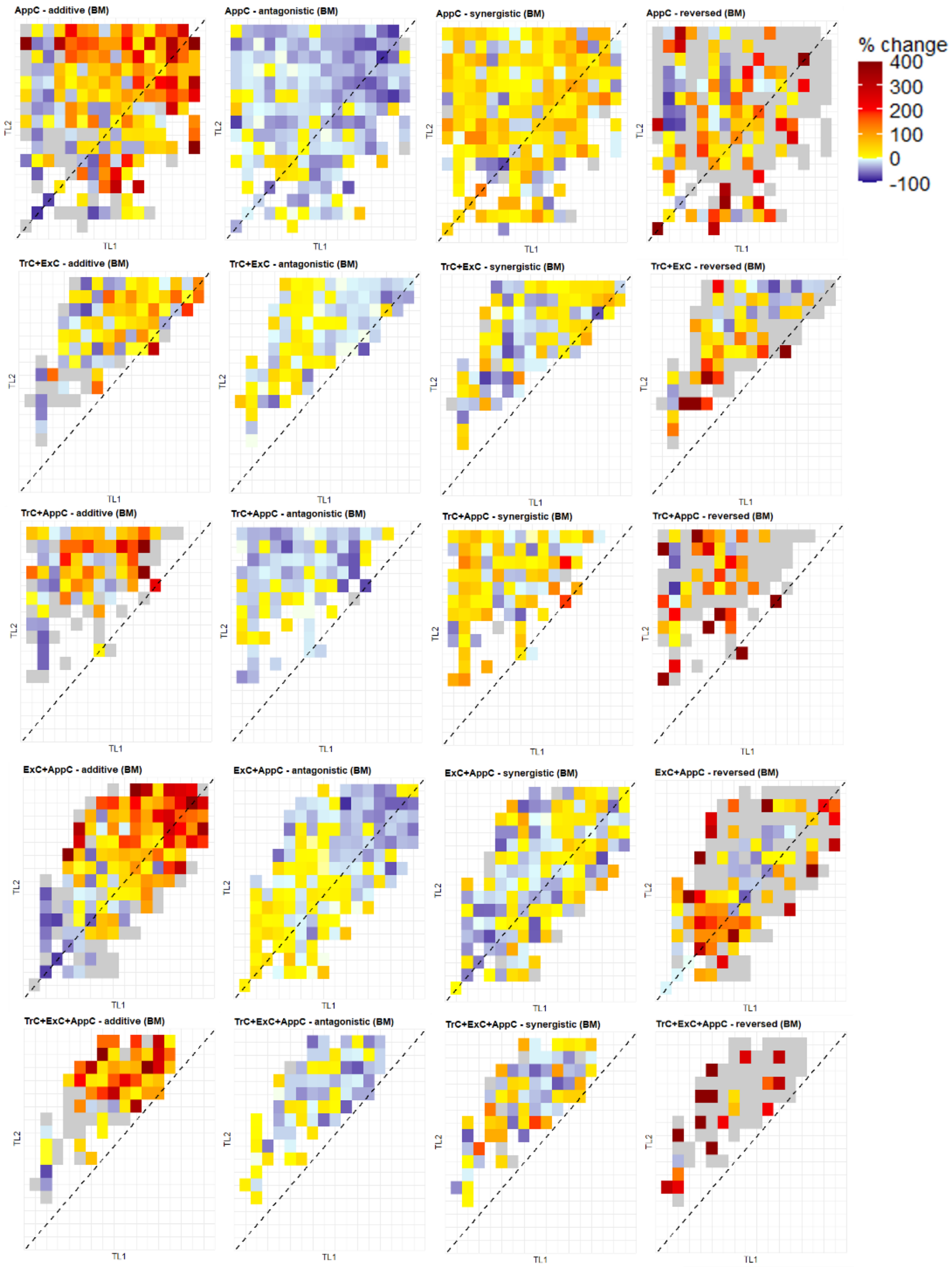


***Figure S6*** *(continued).*
